# TRAIL induces caspase-dependent cleavage of specific components of the clathrin-mediated endocytosis machinery

**DOI:** 10.1101/2024.10.24.619217

**Authors:** David A Lawrence, Scot A Marsters, Cary Austin, Avi Ashkenazi

**Author notes:** Address correspondence to and/or.

## Abstract

It has been reported that caspase activation by the proapoptotic ligand Apo2L/TRAIL disrupts the clathrin-mediated endocytosis machinery (CMEM). To confirm whether TRAIL induces caspase-mediated cleavage of specific CMEM components, we examined the temporal and functional relationship between TRAIL-induced processing of the apoptosis-initiating protease, caspase-8, and the apoptosis-executing protease, caspase-3, *versus* cleavage of specific adaptin (AP) and clathrin heavy chain (CHC) proteins. TRAIL induced time-dependent proteolytic processing of caspase-8 and caspase-3, which coincided with cleavage of AP2α and CHC. The pan caspase inhibitor zVAD-FMK, which blocked caspase processing in response to TRAIL, also prevented the cleavage AP2α. Whereas FADD-deficient or caspase-8-deficient cells showed little or no TRAIL-driven cleavage of either AP2α or CHC, Bax-deficient or caspase-3-deficient cell lines retained AP2α cleavage but failed to cleave CHC in response to TRAIL. The DNA-damaging agent doxorubicin also led to processing of caspase-8 and caspase-3 in conjunction ith cleavage of AP2α and CHC. Together, these results confirm that TRAIL induces caspase-dependent cleavage of AP2α and CHC. Whereas TRAIL-driven cleavage of both AP2α and CHC requires caspase-8, cleavage of CHC, but not of AP2α, requires caspase-3.

**MATERIALS AND METHODS:** 

**Cell lines and cell culture:** All cell lines were obtained from ATCC or as kind gifts from Dr. Bert Vogelstein (HCT116 Bax^-/-^) and Dr. John Blenis (Jurkat I9.2 and E1). Cell lines were cultured with standard media as previously described (1).

**Immunoblot analysis:** Cells were lysed in RIPA lysis buffer (20-188, Millipore) supplemented with Halt protease and phosphatase inhibitor cocktail (ThermoFisher Scientific) and kept on ice for 30 min. Lysates were cleared by centrifugation at 13,600 x g for 15 min at 4 °C, and protein amount was determined by BCA protein assay (ThermoFisher Scientific). Protein was denatured by adding NuPAGE LDS buffer and DTT reducing buffer (Invitrogen) and incubating the samples at 95 °C for 5 min. Equal amounts of denatured protein were loaded into each well of NuPAGE pre-cast gels (Invitrogen), resolved by SDS-PAGE, and electro-transferred to nitrocellulose membranes using the iBLOT2 system (Invitrogen). Membranes were blocked in a 5% nonfat milk solution for 1 hr at room temperature and probed with the corresponding primary antibody at 1:1,000 dilution overnight at 4 °C. This was followed by incubation with the corresponding horseradish peroxidase (HRP)- conjugated secondary antibody at 1:10,000 dilution during 1 hr at room temperature. All secondary antibodies were from Jackson Laboratories. The primary antibodies and secondary HRP-conjugated antibodies are listed respectively in **Table 1** and **Table 2**. β-actin IB was used to verify uniformity of protein loading and electro-transfer.

**Additional reagents:** Recombinant soluble non-tagged human TRAIL (Apo2L.0) and Flag-tagged TRAIL were prepared at Genentech. Flag-tagged TRAIL was crosslinked with anti-Flag M2 antibody at a ligand-to-antibody molar ratio of 1:2.

zVAD-FMK was purchased from R & D Systems, FMK001.

Doxorubicin was purchased from Sigma Aldrich, D1515.

## INTRODUCTION

We have previously reported that the proapoptotic ligand Apo2L/TRAIL (TNFSF10) induces clathrin-mediated endocytosis (CME) of its cognate death receptor, DR5, and that apoptosis activation by TRAIL disrupts the CME pathway (1). Furthermore, inhibition of CME by a dominant-negative mutant of dynamin 1 augmented TRAIL-driven activation of caspase-8 and caspase-3, suggesting that endocytosis of the death-inducing signaling complex assembled by TRAIL dampens proapoptotic signaling (1). A subsequent study provided independent evidence that inhibition of endocytosis by exposure of cells to hyperosmotic glucose enhanced TRAIL-induced activation of caspase-8 and caspase-3 (2). Moreover, a global proteomic study of Jurkat cells indicated that TRAIL induces caspase-mediated cleavage of several components of the CME machinery (CMEM), including CLTA, AP2A1, AP2A2, and DNM2, amongst other caspase-targeted proteins (3). Here we provide additional evidence confirming that TRAIL induces caspase-dependent cleavage of the key CMEM components AP2α and CHC. While AP2α cleavage requires caspase-8 but not caspase-3, CHC cleavage requires both caspases.

**Table 1.**
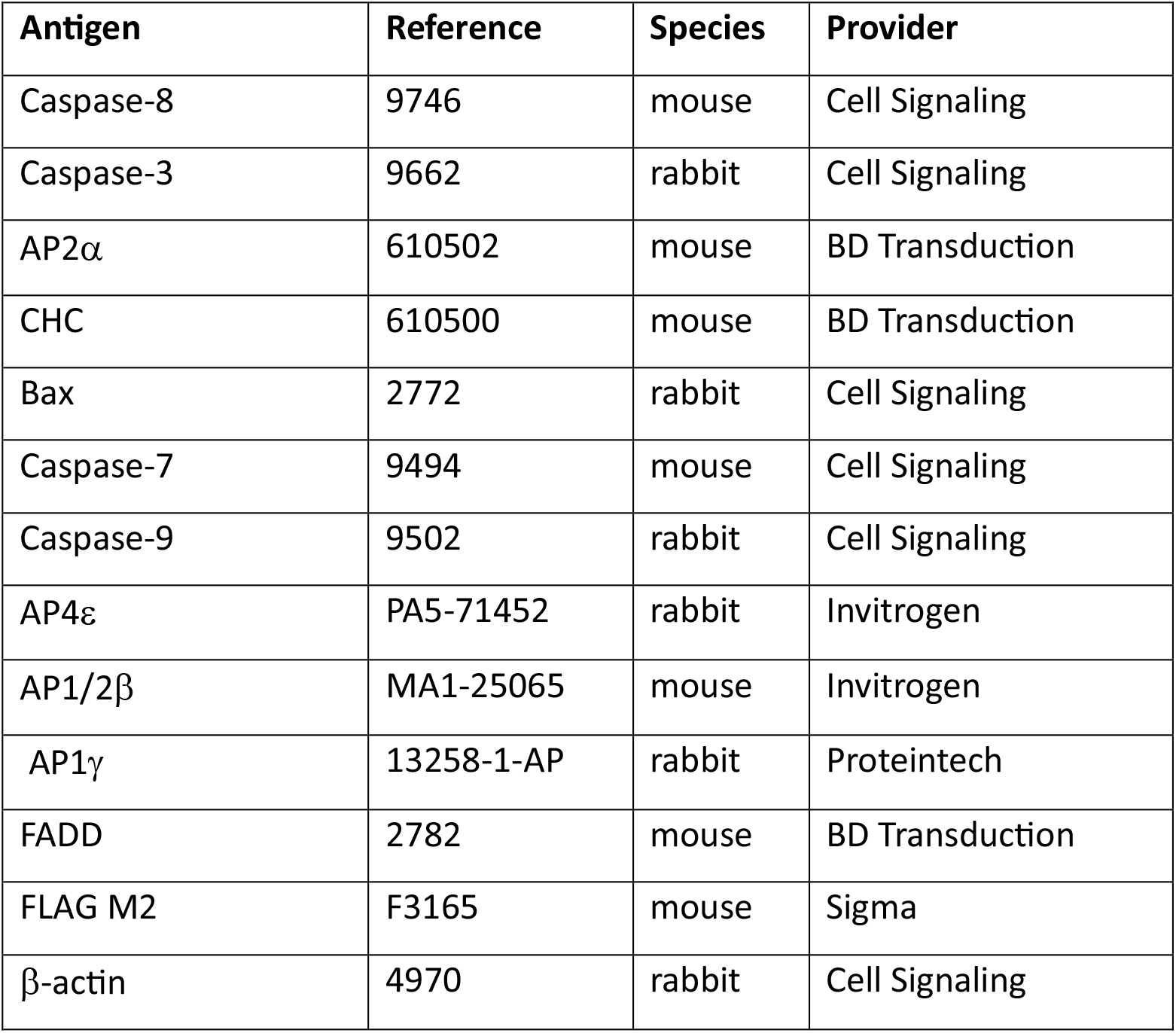
Primary antibodies used for IB analysis.

| **Antigen** | **Reference** | **Species** | **Provider** |
| --- | --- | --- | --- |
| Caspase-8 | 9746 | mouse | Cell Signaling |
| Caspase-3 | 9662 | rabbit | Cell Signaling |
| AP2α | 610502 | mouse | BD Transduction |
| CHC | 610500 | mouse | BD Transduction |
| Bax | 2772 | rabbit | Cell Signaling |
| Caspase-7 | 9494 | mouse | Cell Signaling |
| Caspase-9 | 9502 | rabbit | Cell Signaling |
| AP4ε | PA5-71452 | rabbit | Invitrogen |
| AP1/2β | MA1-25065 | mouse | Invitrogen |
| AP1γ | 13258-1-AP | rabbit | Proteintech |
| FADD | 2782 | mouse | BD Transduction |
| FLAG M2 | F3165 | mouse | Sigma |
| β-actin | 4970 | rabbit | Cell Signaling |

**Table 2.** Horseradish peroxidase conjugated secondary antibodies used for IB analysis.

| **Antigen** | **Reference** | **Provider** |
| --- | --- | --- |
| Rabbit IgG | 711-035-152 | Jackson Labs |
| Mu IgG2b | 1090-05 | Southern Biotech |
| Mu IgG1 | 559626 | BD Transduction |

## RESULTS AND DISCUSSION

We began this study by investigating the temporal relationship between TRAIL-induced processing of the pivotal proapoptotic caspases, caspase-8 (C8) and caspase-3 (C3), and cleavage of specific CMEM components. We first examined the Colo205 cell line, previously shown to be sensitive to TRAIL-induced apoptosis (4, 5). We treated Colo205 cells with TRAIL for specific time durations, ranging from 0 to 120 min; we then collected whole cell lysates and visualized cellular proteins of interest by immunoblot (IB) analysis. TRAIL induced processing of C8, detectable within 2 min of addition, and becoming more complete by 120 min (**Fig. 1a**). TRAIL also promoted processing of C3, detectable by 30 min, and becoming more complete by 120 min (**Fig. 1a**). These proteolytic processing events were accompanied by discernible cleavage of AP2α by 30 min; of AP1/2β by 60 min; and of CHC by 120 min (**Fig. 1a**). In contrast to Colo205 cells, treatment of the TRAIL-resistant cell line HCT8 with TRAIL did not induce appreciable changes in the levels of C8 and C3, nor in those of AP2α and CHC (**Fig. 1b**), demonstrating a lack of CMEM-component cleavage in the absence of caspase processing. Even longer incubation periods of up to 24 hr of HCT8 cells with TRAIL did not result in detectable depletion of AP2α or CHC (**Fig. 1b**), further suggesting that cleavage of these CMEM components requires efficient caspase activation. We next examined an additional TRAIL-sensitive cell line, namely, BJAB. As in Colo205 cells, in BJAB cells TRAIL induced cleavage of AP2α by 30-120 min (**Fig. 1c**). In contrast to AP2α, neither AP1γ nor AP4ε levels were significantly affected, suggesting that specific AP subunits differ in susceptibility to TRAIL-driven cleavage.

**Figure 1.**
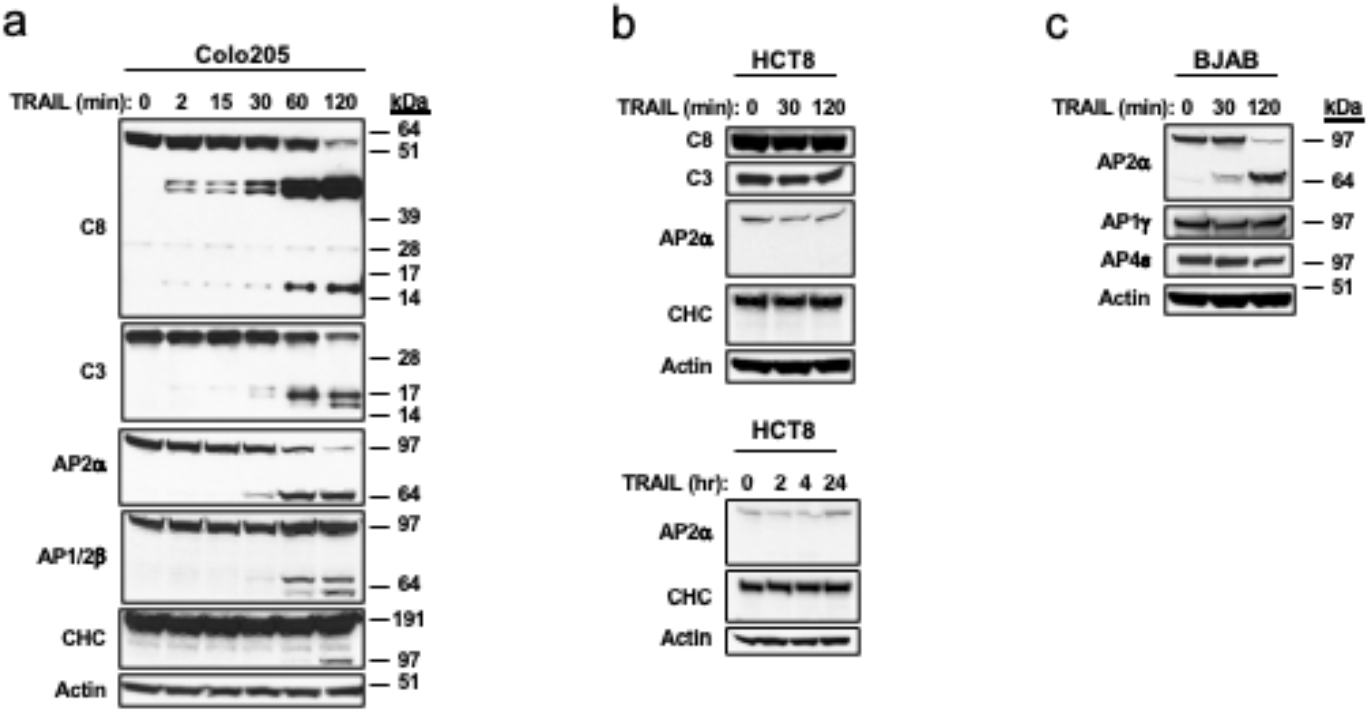
Temporal relationship between TRAIL-induced processing of caspases and cleavage of CMEM components. (**a**) Colo205 cells were treated with recombinant soluble non-tagged TRAIL (0.2 μg/mL) for the indicated time and the noted proteins were analyzed by IB. (**b**) HCT8 cells were treated with recombinant soluble non-tagged TRAIL (0.2 μg/mL) for 0-120 min (top) or 0-24 hr (bottom) and the noted proteins were analyzed by IB. (**c**) BJAB cells were treated with recombinant soluble Flag-tagged TRAIL (1 μg/mL) and anti-Flag antibody (2 μg/mL) for the indicated time and the noted proteins were analyzed by IB.

Thus, TRAIL can induce rapid processing of C8, followed by processing of C3, consistent with its known sequential activation of these proteases (6). Concurrent with a more complete processing of C8 and C3, TRAIL induces sequential cleavage of AP2α, AP1/2β, and CHC. Cleavage of AP2α and CHC occurs in cell lines sensitive to, but not in a cell line resistant to, TRAIL-induced apoptosis. Taken together, these results suggest that the cleavage of CMEM components in response to TRAIL requires effective proapoptotic caspase activation.

To examine the requirement for caspase activation more directly, we leveraged the pan-caspase inhibitor zVAD-fluoromethyl ketone (zVAD-FMK, or zVAD). As expected, treatment of BJAB cells with TRAIL for 4 hr led to processing of C8, C9, and C3 as well as to cleavage of AP2α and CHC (**Fig. 2a**). Also as predicted, zVAD addition blocked C8, C9, and C3 processing completely (**Fig. 2a**), confirming the inhibitors ability to fully prevent caspase activation in this context. Importantly, zVAD also blocked the cleavage of AP2α and CHC (**Fig. 2a**), demonstrating that both events require caspase activation. Consistent with these latter results, in A549 cells zVAD also attenuated the processing of C8, C9 and C3, as well as the cleavage of AP2α (**Fig. 2b**). Next, we used HCT116 cells, in which robust TRAIL-induced C3 activation requires signal amplification downstream to C8 through the intrinsic/mitochondrial apoptosis pathway, engaged via the pro-apoptotic BCL2-family protein Bax (7). We compared a derivative HCT116 cell line harboring a homozygous *Bax* deletion (Bax^-/-^) with the parental HCT116 cell line, which harbors a hemizygous loss of *Bax* (Bax^+/-^). In both cell lines, TRAIL induced a similar processing pattern of C8, albeit diminished somewhat in Bax^+/-^ cells (**Fig. 2 c and d**). TRAIL induced a similar cleavage pattern of AP2α in both cell lines (**Fig. 2 c and d**), indicating that *Bax* is dispensable for TRAIL-induced AP2α cleavage. Given the known requirement of Bax for TRAIL-induced C3 activation in this cell line (7), these data also suggest that AP2α cleavage in response to TRAIL does not require C3 activation. To further assess this, we leveraged the MCF7 cell line, previously documented to be C3 deficient (8). As expected, C3 was not detected in MCF7 cells (**Fig. 2e**). Treatment of MCF7 cells with TRAIL promoted processing of C8 as well as cleavage of AP2α but did not induce cleavage of CHC (**Fig. 2e**). These results show that TRAIL-driven AP2α cleavage proceeds independently of C3, while CHC cleavage requires C3.

**Figure 2.**
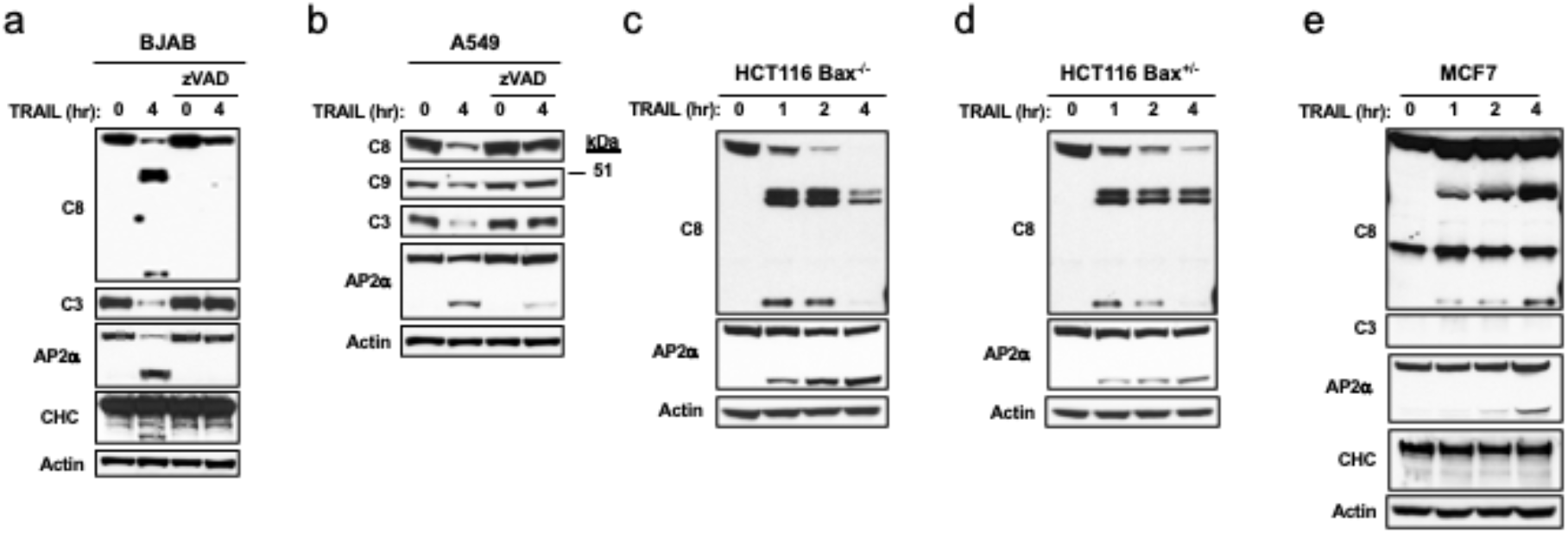
Effect of caspase disruption on TRAIL-induced cleavage of CMEM components. (**a**) BJAB cells were treated with recombinant soluble Flag-tagged TRAIL (1 μg/mL) plus anti-Flag antibody (2 μg/mL) for the indicated time in the absence or presence of zVAD (20 μM) and the noted proteins were analyzed by IB. (**b**) A549 cells were treated and analyzed as in **a**. (**c, d**) HCT116 cell lines harboring homozygous (**c**) or hemizygous (**d**) deletion of Bax were treated with recombinant soluble non-tagged TRAIL (0.2 μg/mL) for the indicated time and the noted proteins were analyzed by IB. (**e**) MCF7 cells were treated and analyzed as in **a**.

Given that AP2α cleavage nevertheless depended on caspase activity–as demonstrated by its inhibition with zVAD–we considered the possibility that AP2α cleavage requires C8, which operates upstream to C3 in the extrinsic apoptotic pathway (6, 9, 10). To investigate this possibility, we used two derivatives of the Jurkat cell line, previously documented to harbor a deficiency in C8 or in the adapter protein FADD, which recruits C8 to death receptors 4 and 5 upon binding to TRAIL (6). As expected, in the parental Jurkat A3 cell line, which expressed FADD and C8, TRAIL induced processing of C8, as well as coincident cleavage of AP2α, and somewhat delayed processing of CHC (**Fig. 3**). In contrast, in the C8-deficient Jurkat variant I9.2, which expressed FADD but not C8, TRAIL failed to induce detectable cleavage of AP2α or new cleavage products of CHC (**Fig. 3**). Moreover, in the FADD-deficient Jurkat variant E1, which expressed C8 but not FADD, TRAIL induced minimal processing of C8, minimal cleavage of AP2α, and no new cleavage products of CHC (**Fig. 3**). It is conceivable that some minor signaling from death receptors to C8 proceeds via alternative adapters such as TRAF2 and/or TRADD (11), which would explain the weak C8 processing and AP2α cleavage still observed in the absence of FADD. Taken together with the data shown in **Fig. 1** and **Fig. 2**, the results in **Fig. 3** indicate that TRAIL-induced AP2α cleavage requires C8 but not C3, whereas CHC cleavage requires both C8 and C3.

**Figure 3.**
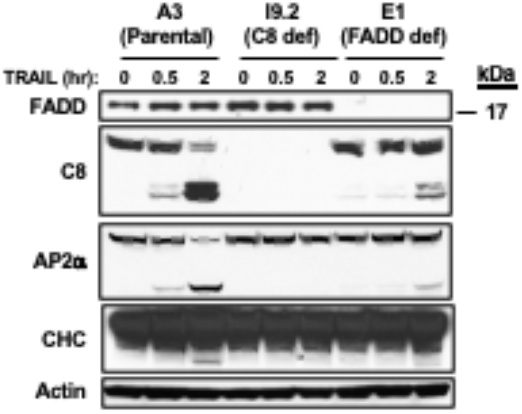
Effect of deficiency in caspase-8 or FADD on cleavage of CMEM components in response to TRAIL. Parental (A3), caspse-8-deficient (I9.2) or FADD-deficient (E1) Jurkat cell lines were treated with recombinant soluble Flag-tagged TRAIL (1 μg/mL) plus anti-Flag antibody (2 μg/mL) for the indicated time and the indicated proteins were analyzed by IB.

To examine whether a different type of proapoptotic driver besides TRAIL could also induce cleavage of CME components, we investigated the effect of the DNA-damaging chemotherapeutic agent, doxorubicin. Treatment of BJAB or Colo205 cells with doxorubicin for 24 hr lead to processing of C8, C3, C7, as well as to cleavage of AP2α and CHC (**Fig. 4**). Induced cleavage of AP1/2β was more clearly detected in BJAB than Colo205 cells. Of note, doxorubicin promoted AP2α and CHC cleavage by 24 hr, while TRAIL provoked these changes already within 30-120 min of addition (**Fig. 1-3**). This temporal difference may be because doxorubicin induces caspase activation indirectly, as a secondary consequence of DNA damage, whereas TRAIL engages the apoptotic pathway more directly, through its cognate death receptors (11). An additional factor that may contribute to kinetic differences between these stimuli is that C3 likely acts upstream to C8 in response to doxorubicin, whereas C8 acts upstream to C3 in response to TRAIL (12). Regardless, these results show that caspase activation by a proapoptotic ligand or a DNA-damaging agent can promote proteolytic cleavage of specific CMEM components, thereby likely altering the cell’s capacity for CME-based internalization of proteins and protein complexes.

**Figure 4.**
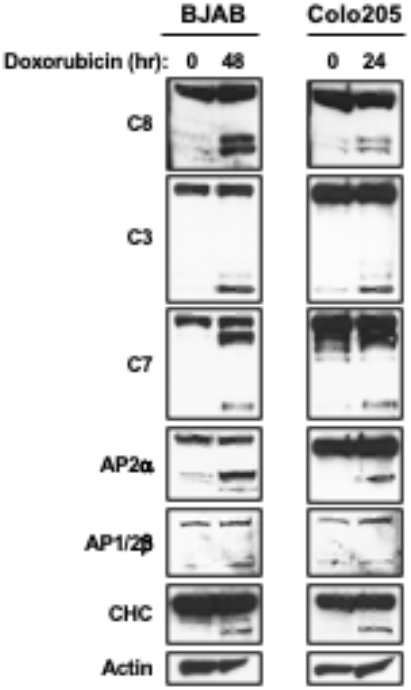
Doxorubicin induces caspase processing and cleavage of CMEM components. BJAB or Colo205 cells were treated with doxorubicin (200 nM) for the indicated time and the noted proteins were analyzed by IB.

## REFERENCES

1. Austin, C et al./person-group>., Proc. Natl. Acad. Sci. USA 103 (27), 10283–10288 (2006).

2. Kohlhaas et al., J. Biol. Chem. 282, 12831–12841 (2007).

3. Stoehr et al., Mol. Cell. Proteomics 12, 1436–1450 (2013).

4. Ashkenazi et al J. Clin. Invest. 104 (2), 155–162 (1999).

5. Wagner et al., Nature Med. 13 (9), 1070–1077 (2007).

6. Kischkel et al., Immunity 12 (6), 611–620 (2000).

7. LeBlanc et al., Nature Med. 8 (3), 274–281 (2002).

8. Jänicke, R.U. Breast Cancer Res. Treat. 117, 219–221 (2009).

9. Ashkenazi and Dixit, Science 281 (5381), 1305–1308 (1998).

10. Ashkenazi and Dixit, Curr. Opin. Cell Biol. 11 (2), 255–260 (1999).

11. Ashkenazi and Salvesen, Annu. Rev. Cell. Dev. Biol. 30 (1), 337–356 (2013).

12. Salvesen and Ashkenazi Cell 147 (5), 1197 (2011).

